# SIEGA the integrated genomic surveillance system of Andalusia, a One Health regional resource connected with the clinic

**DOI:** 10.1101/2024.02.29.582716

**Authors:** Carlos S. Casimiro-Soriguer, Javier Pérez-Florido, Enrique A. Robles, María Lara, Andrea Aguado, Manuel A. Rodríguez Iglesias, José A. Lepe, Federico García, Mónica Pérez-Alegre, Eloísa Andújar, Victoria E. Jiménez, Lola P. Camino, Nicola Loruso, Ulises Ameyugo, Isabel María Vazquez, Carlota M. Lozano, J. Alberto Chaves, Joaquin Dopazo

## Abstract

The One Health approach, recognizing the interconnectedness of human, animal, and environmental health, has gained significance amid emerging zoonotic diseases and antibiotic resistance concerns. This paper aims to demonstrate the utility of a collaborative tool, the SIEGA, for monitoring infectious diseases across domains, fostering a comprehensive understanding of disease dynamics and risk factors, highlighting the pivotal role of One Health surveillance systems.

Raw whole-genome sequencing is processed through different species-specific open software that additionally reports the presence of genes associated to anti-microbial resistances and virulence. The SIEGA application is a Laboratory Information Management System, that allows customizing reports, detect transmission chains, and promptly alert on alarming genetic similarities.

The SIEGA initiative has successfully accumulated a comprehensive collection of more than 1900 bacterial genomes, including *Salmonella enterica*, *Listeria monocytogenes*, *Campylobacter jejuni*, *Escherichia coli*, *Yersinia enterocolitica* and *Legionella pneumophila*, showcasing its potential in monitoring pathogen transmission, resistance patterns, and virulence factors. SIEGA enables customizable reports and prompt detection of transmission chains, highlighting its contribution to enhancing vigilance and response capabilities.

Here we show the potential of genomics in One Health surveillance when supported by an appropriate bioinformatic tool. By facilitating precise disease control strategies and antimicrobial resistance management, SIEGA enhances global health security and reduces the burden of infectious diseases. The integration of health data from humans, animals, and the environment, coupled with advanced genomics, underscores the importance of a holistic One Health approach in mitigating health threats.

## Introduction

The One Health approach emphasizes the interconnectedness of human, animal, and environmental health, and has gained relevance in recent years due to the emergence of zoonotic diseases and increasing antibiotic resistance ^1^. The importance of One Health in epidemiological surveillance lies in its capacity to monitor and control infectious diseases across different domains, enabling a comprehensive understanding of disease dynamics and risk factors ^2^. By integrating human, animal, and environmental health data, One Health surveillance systems facilitate early detection and response to health threats, thereby enhancing global health security and reducing the burden of disease ^3^. Furthermore, this integrated approach fosters multidisciplinary collaborations among stakeholders to develop and implement coordinated strategies aimed at disease prevention and control ^4^. Recently, the conventional process of serotyping through serology has been undergoing a gradual transformation, with molecular typing methods such as multi-locus sequence-based typing (MLST) ^5^ increasingly complementing or replacing traditional methods. However, these techniques do not possess the necessary discriminatory power to differentiate between closely related strains ^6^, which limits its application in many epidemiological studies. In recent years, High-throughput sequencing (HTS) technologies have revolutionized the field, enabling rapid and cost-effective analyses of complete genomes ^7^. Whole-genome sequences (WGS) offer an unparalleled level of discrimination among genetically related isolates, allowing for the exploration of compelling questions such as accurate phylogenetic and phylogenomic analyses ^8^, as well as the examination of serotype- or subtype-determining genes ^9^. Thus, the use of sequence-derived typing is gaining acceptance and is being employed for source attribution and epidemiological surveillance ^10^.

Accordingly, different analytical strategies have been developed ^11–13^, and numerous studies have demonstrated the potential of WGS in epidemiological investigations ^14–18^. This advancement in genomic sequencing is revolutionizing the way in which the epidemiology of various pathogens is approached and assessed and, actually, the European Centre for Disease Prevention and Control (ECDC) ^19^ and the World Health Organizarion ^20^ have recommended the use of WGS as the gold standard methodology for surveillance of bacterial pathogens. One interesting aspect of WGS is its ability to provide not only precise typing but also the opportunity to conduct additional analyses beyond routine surveillance, such as assessing the existence of genetic factors related to antimicrobial resistance and virulence, which holds great significance in the context of the One Health approach ^21^. This approach is rapidly emerging as the primary framework for monitoring and managing antimicrobial resistance, establishing a compelling connection between genomics and comprehensive control strategies.

Andalusia, with a population of 8.5 million inhabitants, is the third largest region in Europe and has the size of a medium-sized European country, like Switzerland or Austria. Here we present a region-wide genomic surveillance system, SIEGA (acronym for “Integrated system of genomic epidemiology in Andalusia” in Spanish). As of July 2023, SIEGA contains a total of 1906 bacterial genomes corresponding to 670 *Salmonella enterica*, 688 *Listeria monocytogenes*, 276 *Campylobacter jejuni*, 191 *Escherichia coli,* 23 *Yersinia enterocolitica* and 58 *Legionella pneumophila*, collected at the different provinces of Andalusia in food products, factories, farms, water systems and human clinical samples (although these numbers change rapidly due to the continuous increase of samples). SIEGA contains information on the origin of the samples, their typing and resistance and virulence genes. It consist of two modules, one of them is public and contains descriptive information on the samples in the conventional NEXSTRAIN representation ^22^, and the other one, the SIEGA management data system, is a private Laboratory Information Management System (LIMS) for the use of the personnel of public health and the clinicians involved in the surveillance program. The LIMS allows users to build customized reports on the isolates, provides tools for detecting transmission chains and implements an automated alert system that promptly signals whenever a newly detected bacterium exhibits a predetermined genetic similarity to any existing database entry, enhancing vigilance and response capabilities. Some statistics on the isolates as well as a more detailed study on the relationships between them and the distribution of resistances across samples is provided.

## Results

### The SIEGA circuit

On May 15, 2020, the Andalusian Local Ministry of Health entrusted the Progress and Health Foundation to initiate a program aimed at monitoring pathogens through whole genome sequencing and bioinformatic analysis of a specific set of bacteria with significant implications in public health. ^23^. This marked the inception of the SIEGA initiative.

SIEGA has been conceived as an integrated system aimed at incorporating samples from all relevant sources, such as clinical, food, environmental, industry or primary production samples. In its early stages, the initial samples were sequenced in a privately contracted laboratory. Subsequently, the Regional Ministry of Health and Consumer Affairs designed a genomic sequencing circuit aimed at facilitating this integration, which was included in Instruction 130/2019 ^24^ on the treatment and sequencing of biological agents isolates in Andalusia with a focus on health protection, inspired in The Transformation of Reference Microbiology Methods and Surveillance for Salmonella With the Use of Whole Genome Sequencing in England and Wales ^25^. Figure 1 sketches the general operating layout of the circuit.

**Figure 1.**
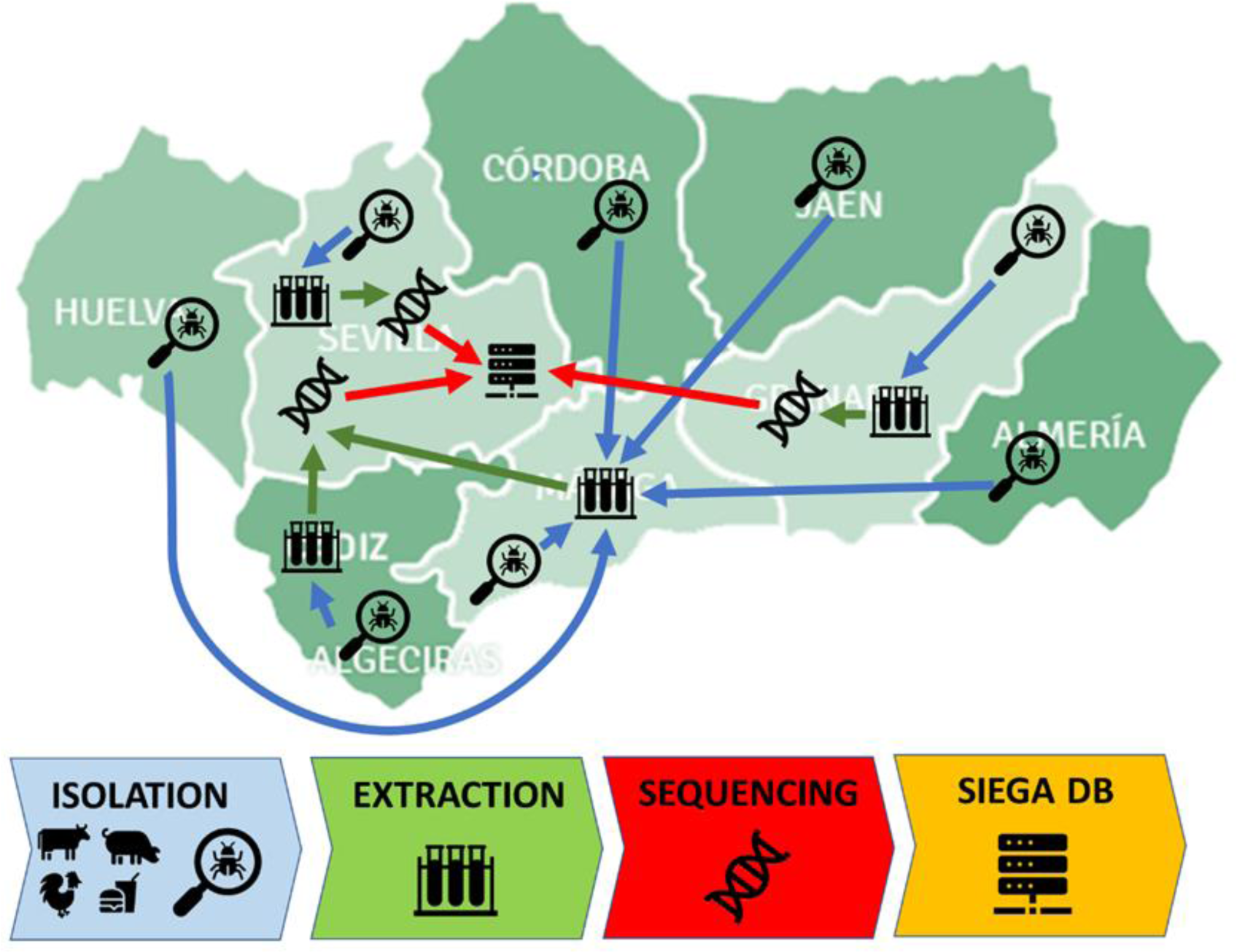
The SIEGA circuit. The different provinces that collect samples send them for extraction of DNA to the different reference laboratories, which is subsequently sent to the sequencing facilities and finally the resulting genomic data is uploaded in the central SIEGA data management system.

Any laboratory that isolates any strain of the pathogens of interest can send it, under the conditions set out in that instruction and accompanied by basic metadata, to the Public Health Laboratory of Málaga, under the Regional Ministry of Health and Consumer Affairs, where DNA extraction is carried out. Once the DNA is extracted, it is preserved in freezing conditions and sent to the sequencing laboratory at the Andalusian Molecular Biology and Regenerative Medicine Center (CABIMER).

Additionally, other collaborating centers have been incorporated, which have either provided DNA extracts (University Hospital Puerta del Mar in Cádiz) or directly provided sequences (University Hospital Virgen del Rocío or University Hospital San Cecilio).

Non-clinical origin samples primarily stem from official veterinary control activities in primary production, such as the implementation of the Annual National Program for the Control of Certain Serotypes of Salmonella in meat chickens of the species *Gallus gallus*. They also originate from Official Control Services in stages subsequent to primary production, which are included in the National Official Control Plan for the Food Chain. This was agreed upon in coordination meetings between the competent authorities. Additionally, other samples from investigations conducted within the framework of the management of foodborne outbreaks are included. All these samples are analyzed in designated official laboratories in accordance with the provisions of Article 37 of Regulation 2017/625, which, in the case of Andalusia, follow the aforementioned Instruction 130/2019.

Clinical origin samples are obtained in the exercise of the healthcare function of the referring centers on a voluntary basis, also applying the aforementioned Instruction 130/2019.

### The public SIEGA webpage

As previously mentioned, SIEGA has a public website ^22^ where a detailed description of the circuit and updated information on the species under surveillance is available for the general public. Having a public webpage for a project of genomic surveillance of pathogens is vital for promoting transparency, disseminating knowledge, fostering collaboration, and bridging the gap between the scientific community and the general public. By providing open access to information on the surveillance achievements, SIEGA encourages participation from diverse stakeholders, and creates a more knowledgeable and engaged society in the ongoing fight against infectious diseases. SIEGA offers access to Nextstrain Auspice ^26^ viewers for the different species under surveillance: *Salmonella enterica, Listeria monocytogenes, Campylobacter jejuni, Escherichia coli, Legionella pneumophila* and *Yersinia enterocolitica* (see Figure 2). In the viewers, the public can explore the relationships among the samples sequenced in the region and other international samples of reference. It is also possible to locate in a map the geographical origin of the different isolates, including the animated options available in the interface, which emulate in a very visual way the transmission of the samples over time.

**Figure 2.**
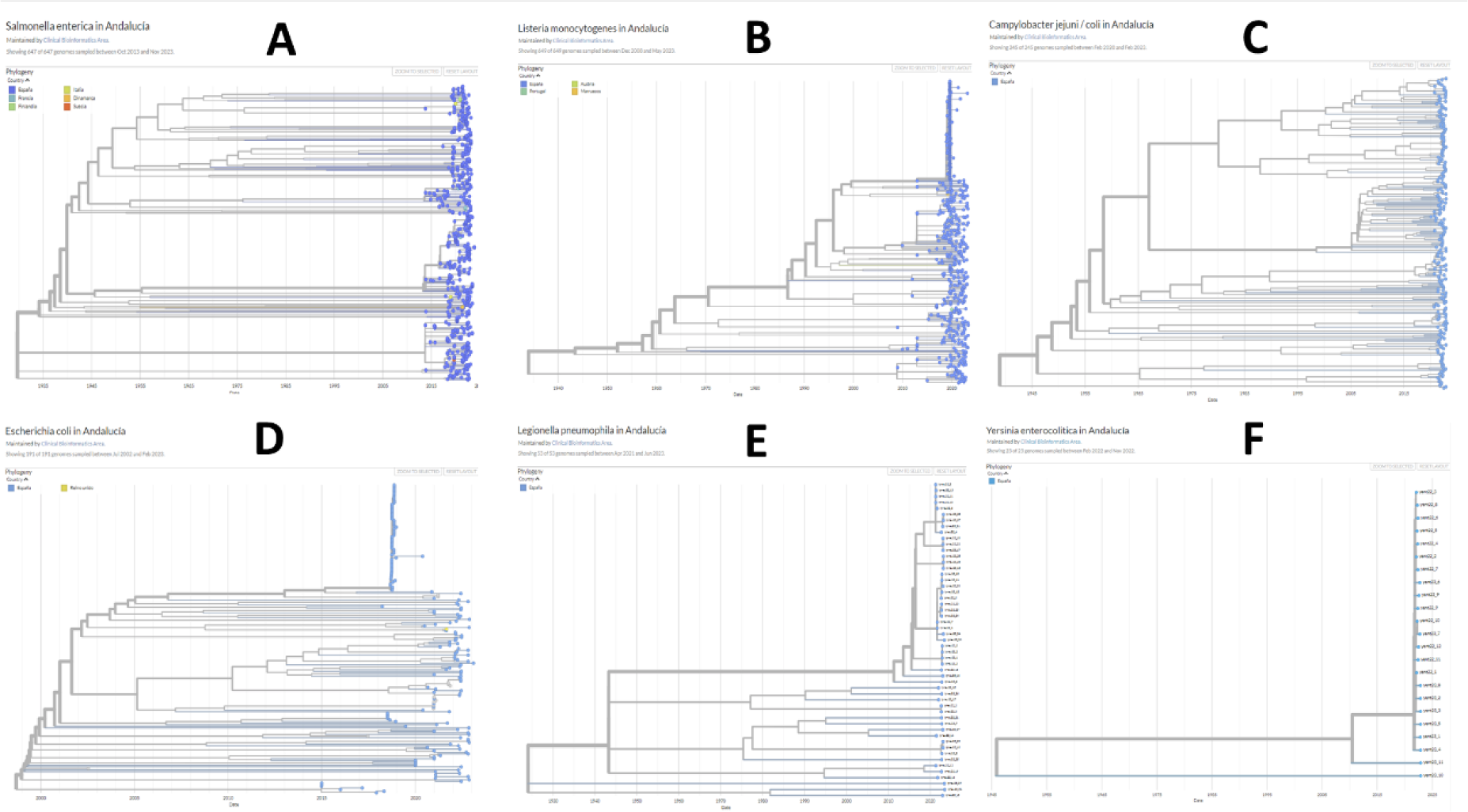
Phylogenies from Nextstrain viewers for: A) *Salmonella enterica*, B) *Listeria monocytogenes*, C) *Campylobacter jejuni*, D) *Escherichia coli*, E) *Legionella pneumophila* and F) *Yersinia enterocolitica*.

### The SIEGA data management system

The SIEGA data management system serves as a private LIMS designed for utilization by personnel in public health and participating clinicians of the surveillance program. SIEGA facilitates the seamless uploading of raw sequencing data and orchestrates automated processing, including quality control assessments. Through this platform, users can generate tailored reports concerning the isolates, explore potential transmission chains, and deploy an automated alert mechanism that promptly signals any genetic similarity between newly identified bacteria and entries within the existing database. This system bolsters vigilance and response capabilities. Furthermore, SIEGA furnishes statistical insights into the isolates, along with a comprehensive exploration of their interrelationships and the distribution of resistances across samples.

Within the SIEGA interface, each organism has five distinct subsections to facilitate comprehensive analysis that include sample status, metadata, control results and Flexible Table, a wizard to combine metadata for complex representations (see Table 1).

**Table 1.** SIEGA features and description.

| Feature | Description |
| --- | --- |
| Interface | With different subsections to facilitate the analysis:<br>• **Sample Status:** information on each sample, including the quality control status<br>• **Metadata:** information needed for sample and study traceability.<br>• **Application control:** information on the status and results of the pipeline of analysis<br>• **Results:** detailed overview of the results obtained for any sample<br>• **Flexible Table:** allows merging of any variety of result tables, metadata, or statuses for complex representation of results |
| User management | Access permission structure allowing users and group with many options to share data across users and groups |
| Data upload | • Comma-separated values (CSV) file for metadata<br>• Fastq R1 and R2 files for each sample |
| Data processing and quality control | Quality control information on the result of each application is reported (green: good, yellow: fair, red: poor). |
| Data traceability | Each sequenced sample has a distinctive identifier, used for associating relevant metadata |
| Automatic reporting | A report is generated for each sample including:<br>• MLST<br>• cgMLST<br>• serotyping outcomes<br>• antimicrobial resistance genes<br>• virulence genes<br>• plasmids<br>• a phylogenetic tree with the most related samples |
| Customized phylogenetic analysis | All existing tables can be combined using a wizard (flexible table) that enables the creation of combined tables with relevant data for each case, from which phylogenetic trees can be generated |
| Alert system | Automatic alerts based on various criteria can be established:<br>• allelic distances<br>• specific sequence type match<br>• serotype predicted<br>• presence of antimicrobial resistance genes<br>• presence of virulence genes |

The entire SIEGA data management system has been designed to free users of the need to have in-house expertise in genomic data management and resources to store such data. Simultaneously, the system offers a centralized database of all genomes sampled, allowing optimal exploitation of the results and the correct implementation of one health surveillance. It provides a convenient user permission structure to allows data sharing and collaborative work (if desired), detailed quality control for the standard pipelines used for data processing and accurate data traceability. Detailed reports on each sample, that include MLST, cgMLST, serotyping outcomes, antimicrobial resistance and virulence genes, plasmids and a phylogenetic tree with the most related samples are provided. Additionally, customized phylogenetic analysis can be carried out in an easy and intuitive way. Finally, one of the most interesting features is the automatic alert system, that automatically warns when a new sample is introduced that meets some configurable criteria of genetic distance, serotype. Presence of antimicrobial or virulence genes, etc.

### SIEGA interface

Within the SIEGA interface, each organism has five distinct subsections to facilitate comprehensive analysis:

**Sample Status**: This section contains information on the sample, including the quality control status (traffic light system), the date of incorporation into the SIEGA database, and the initiation date of the pipeline, among other details.

**Metadata**: This section contains crucial information needed for traceability, epidemiological studies, and preliminary sample processing, fostering a robust framework for data management.

**Application Control**: Here, users have information on the pipeline of applications implementing the different algorithms that have been applied to process each sample and verify their successful completion.

**Results**: Hare users can access to the most pertinent outcomes derived from each application in the genomic pipeline, providing them with a detailed overview of the mains results obtained for any sample.

**Flexible Table**: This feature enables the merging of any variety of result tables, metadata, or statuses. It further allows for the visualization of merged data on the phylogenetic tree for each organism or selected subgroups of samples.

The entire SIEGA data management system has been designed to free users of the need to have in-house expertise in genomic data management and resources to store such data. Simultaneously, the system offers a centralized database of all genomes sampled, allowing optimal exploitation of the results and the correct implementation of one health surveillance.

### User management

A practical data access permission structure has been implemented in the SIEGA data management system. Each authorized SIEGA user is allocated to a specific group and can only view and work with samples within their own group, publicly shared samples from other groups, and samples shared with their group by others. Additionally, their group membership determines their access to various metadata sets. This strategy ensures that sensitive metadata is visible only to specific groups.

Within each group, there is an assigned administrator who holds the necessary permissions to upload new samples, share samples with other groups, and make samples universally available to all groups. This allows each group to maintain control over the samples they upload and their visibility to other groups, whilst safeguarding sensitive data.

There are only two groups with special permissions: i) Andalusia’s Epidemiological Surveillance System and ii) Health Protection. In the event of an outbreak involving any organism, these groups have the necessary access to view crucial data for taking action and containing the situation.

### Data upload

Data uploading is streamlined into a user-friendly three-step wizard. In the initial step, users download a Comma-separated values (CSV) file containing metadata for database incorporation. Next, the second step entails uploading this completed CSV metadata file and confirming successful data integration. Finally, in the third step, users upload the fastq R1 and R2 files for each sample, ensuring their accuracy, absence in the database, and precise name matching as specified in the metadata.

### Data processing and quality control

Upon the upload of the fastq files, the genomic analysis pipeline is sequentially initiated for each sample. As quality control data for each application is generated, it is represented in the sample status table using a traffic light system (green: good, yellow: fair, red: poor). This provides a rapid evaluation of sample quality. Moving the cursor over the traffic light icon reveals a pop-up window with specific details, including the application utilized and quality metrics. Samples with any quality parameter marked as red are omitted from subsequent analyses.

### Data traceability

Each sequenced sample within SIEGA is allocated a distinctive identifier, prominently displayed in the viewer interface. This identifier serves as a hub for associating relevant metadata. Depending on the sample type, specific identifiers are appended. For instance, food and environmental samples receive an “Albega” number, which links to establishment details, product type, date, results, associated inspector, and more. Livestock samples are tagged with the “livestock farm” number, encompassing farm specifics, animal type, sampling area, date, etc. In clinical samples, the NUHSA (Unique Health History Number of Andalusia) is utilized, that connects with a comprehensive personal health history, incorporating age, sex, pathologies, type of analyzed sample, and additional data, establishing a robust One Health framework for sample origin identification.

### Automatic reporting

For each sample, a report is generated, presenting the most relevant genomic analysis results. It includes essential quality metrics, sample characterization with MLST, cgMLST, and serotyping outcomes, as well as information about antimicrobial resistance genes, virulence genes, plasmids, the phylogenetic tree encompassing samples within 50 Single Nucleotide Polymorphism (SNPs) of genetic distance, and the list of samples within 10 allelic differences. Users can access this information in HTML format and also download it as a PDF.

### Customized phylogenetic analysis

In the flexible table section, all existing tables in the web application can be combined using a wizard based on the unique sample ID. This enables the creation of tables with relevant data for each case, and once created, phylogenetic trees can be generated from the selected samples. Each sample will be color-coded based on the data present in the table, using both GrapeTree ^27^ and Taxonium ^28^.

### Alert system

One of the most interesting functionalities of the SIEGA application is the capability to establish automatic alerts based on various criteria. These alerts can be tailored to the user’s preferences by utilizing both metadata and the outcomes of genomic analysis (i.e. allelic distances, specific sequence type match, serotype predicted, presence of antimicrobial resistance or virulence genes) of any new bacterial genome sequence introduced in the database to the rest of sequences already present. The duration of these alerts can be adjusted, incorporating either temporal limits or execution thresholds. Moreover, the system facilitates the sharing of alerts with other user groups. Furthermore, the application enables the direct transformation of a database search filter into the criteria for configuring an alert.

### Advantages of SIEGA

The overarching goal of the SIEGA initiative, and particularly its SIEGA data management system, is to streamline the implementation of epidemiological surveillance, especially at the regional level or within large communities engaged in the One Health approach. It offers a continuously upgraded environment for managing genomic data, eliminating the need for end-users to establish their own bioinformatics teams for data processing and interpretation, as well as to invest in computer infrastructure for data storage and analysis. Furthermore, because data undergo uniform processing through a regularly updated pipelines adhering to international analysis standards, the results are consistent and can be readily compared. This democratizes genomic surveillance by involving all necessary stakeholders, as the SIEGA platform provides the essential resources for both data processing and interpretation.

### Current users

Microbiology laboratories across different hospitals in Andalusia are actively utilizing SIEGA. In addition, the “Sistema de Vigilancia Epidemiológica de Andalucía” (SVEA) from the “Junta de Andalucía” and professionals in the field of Health Protection are also users of SIEGA. In adherence to the One Health principle, individuals involved in both animal health management and laboratory aspects have been incorporated as users of SIEGA.

### Molecular epidemiology of isolates

#### Salmonella enterica

The SIEGA encompasses a dataset comprising 670 whole genome sequences of *Salmonella enterica*, which were sequenced from June 2020 to July 2023 using samples collected between 2013 and 2023. Within this dataset, 42.54% (285) of the samples were sourced from clinical origins, 34.63% from the food-related sector, and 21.34% from livestock sources. A total of 448 distinct Sequence Types (STs) were identified and categorized into clonal complexes (CCs). The prevailing ST, ST 309694 (corresponding to clonal complex ST-71), was encountered on 26 occasions. Additionally, 83 strains exhibited concurrence with more than one ST. 6 STs (ST-67337, ST-138467, ST-197094, ST-207307, ST-247937, and ST-320298) were found cross-wide clinical, food-related, and livestock-origin samples. Similarly, 18 STs were identified in both clinical and food samples, and 7 STs were identified in clinical and livestock-origin samples

#### Listeria monocytogenes

The SIEGA includes a dataset comprising 678 whole genome sequences of *Listeria monocytogenes*, which were sequenced from June 2019 to July 2023. Within this dataset, 69.61% (472) of the samples were sourced from clinical origins, including all the samples from Andalusia sequenced by the Neisseria, Listeria and Bordetella Unit of the National Centre for Microbiology in Spain while investigating the Listeriosis outbreak caused by contaminated stuffed pork in Spain in 2019 ^29^, and 30.38% (206) from food origin. A total of 248 distinct STs were identified and categorized into CCs. The prevailing ST, ST 29514 (corresponding to clonal complex ST-388), was encountered on 210 occasions.

#### Campylobacter spp

The SIEGA contains 276 whole genome sequences of *Campylobacter*, received between December 2020 and June 2023, corresponding to both *C. jejun*i and *C. coli*. Most of the sequences have been obtained from human clinical strains from two reference hospitals in Cádiz and Seville. There are some STs, grouped into CCs, which have been detected with greater frequency in clinical samples. From ST-16294 (corresponding to the clonal complex ST-206) 12 sequences have been obtained, with the interest of being detected from 2020 to 2023 and in a scattered way, with 8 isolates in Cádiz and 4 in Seville. Their identical virulome and resistome profiles have been recovered from these sequences, using the tools described, particularly ABRicate ^30^ on VFDB ^31^ and CARD ^32^, databases. Another frequent STs have been ST-12550 (ST-573CC) and ST-18855 (ST-52CC).

#### Escherichia coli

The SIEGA includes 121 whole genome sequences of *Escherichia coli*, 44 of them downloaded from the EnteroBase website ^33^ for reference and 77 of them collected between November 2021 and May 2023. To date, most of the sequences (72) have been obtained from food samples taken at retail level on behalf of the monitoring programme of anti-microbial resistance (AMR) according to the provisions of the Commission Implementing Decision (EU) 2020/1729 ^34^ implemented in Andalusia. The human clinical strains come from two reference hospitals in Cádiz and Seville. There are STs, grouped into CCs, which have been detected with greater frequency in food samples. The prevailing ST, ST 169652 (corresponding to clonal complex ST-10) was encountered in 6 occasion all from food samples. Additionally, other 4 strains grouped in ST142026 (CC 155) and 3 strains exhibited concurrence with ST 191979 (CC 162) or 60064 (CC 93). To the date, no shared CCs have been detected in the food and human clinical origin samples.

#### Yersinia enterocolitica

The SIEGA encompasses 23 whole genome sequences of *Yersinia enterocolitica*, received in the year 2022, sampled between February and November. To date 21 (91.3%) of the sequences have been obtained from clinical samples from one of the reference hospitals, the Hospital Virgen del Rocio in Seville, and 2 were obtained from food samples. There are some STs, grouped into CCs, which have been detected with greater frequency in these clinical samples. The prevailing ST, ST-1574 (corresponding to clonal complex ST-135), was encountered on 4 occasions. Additionally, 3 strains exhibited concurrence with ST-52 and 2 strains grouped into ST-1716 (corresponding also to clonal complex ST-135). Figure 3 depicts the genetic relationships between all the *Yersinia enterocolitica* samples.

**Figure 3.**
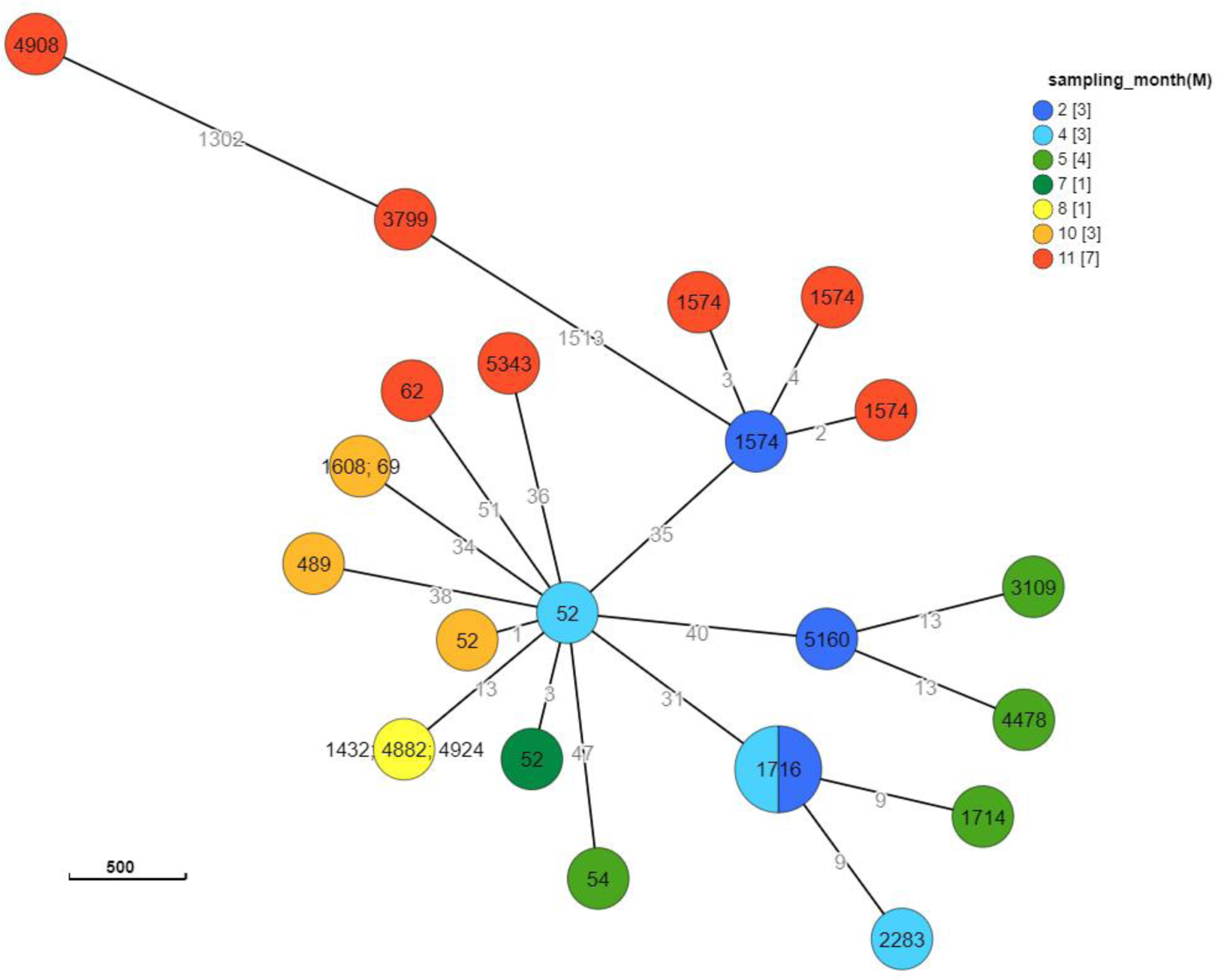
*Y. enterocolitica* GrapeTree representation, generated within the SIEGA application. Node labels represent ST (in some cases an ambiguous ST assignation occurred and more than one number is displayed) and node color correspond to the sampling month (a warm gradient has been used to better display the time scale). Numbers in the branches correspond to the allelic distances among nodes. The GrapeTree representation provides an intuitive visualization of the temporal scale of sampling and the genetic similarities among the samples. Using different labels from the metadata and the results tables, it is possible to obtain visual representations of many aspects of the epidemiology of the selected samples.

#### Legionella pneumophila

Legionella data stored in SIEGA, includes the comparative analyses of 58 *Legionella pneumophila* isolates during 2021-2023. Of these, 12 isolates corresponded to clinical isolates and 46 to environmental isolates. Clonal relation between the Isolates was determined by cgMLST. This scheme classified the isolates into 18 ST (sequence type). The most abundant being ST 293 (20 isolates, 34.5%) and 180F (11 isolates, 18.9%). Four ST (293, 427, 489 and 180F) were present in both clinical and environmental isolates. In addition, we identified 3 STs (95, 98, and 524F) in clinical isolates that are not associated with environmental origin, suggesting that they derived from unrecognized sources.

### Analysis of antimicrobial resistance

In the EU, the new legislation related to the harmonized monitoring and reporting of AMR from 2021 ^35^ authorized whole genome sequencing as an alternative method to supplementary phenotypic testing of *Salmonella* and *E. coli* in certain conditions. The SIEGA allows the monitoring of the presence of resistance genes in the different microorganisms facilitating the tracking of the dissemination or emergence of AMR throughout the food chain under a One Health approach. For example, the presence of AMR genes in the population of Salmonella included in the SIEGA can be analyzed, categorized by antimicrobial classes (Figure 4). This analysis reveals that 52.7% (347) exhibit resistance genes to only 1 group of antimicrobials, while 15% (99) carry resistance genes to two distinct classes of antimicrobials. In contrast, 32.2% (212) demonstrate resistance genes to 3 or more classes of antimicrobials. In a similar manner, this could be carried out with the other microorganisms hosted in the database, or further analysis could be conducted by delving into the multiple variables, for instance, this could involve monitoring the emergence of *Salmonella* strains harboring colistin resistance genes, the occurrence of *Salmonella* strains harboring resistance genes to fluoroquinolones and third-generation cephalosporins or monitoring the presence of resistance genes to Critically Important Antibiotics (CIAs) in the database. The Figure 4 represents a summary of the observed frequency of potential multi-resistances, represented as the number of different AMR genes corresponding to different antimicrobial classes harbored by each individual sample.

**Figure 4.**
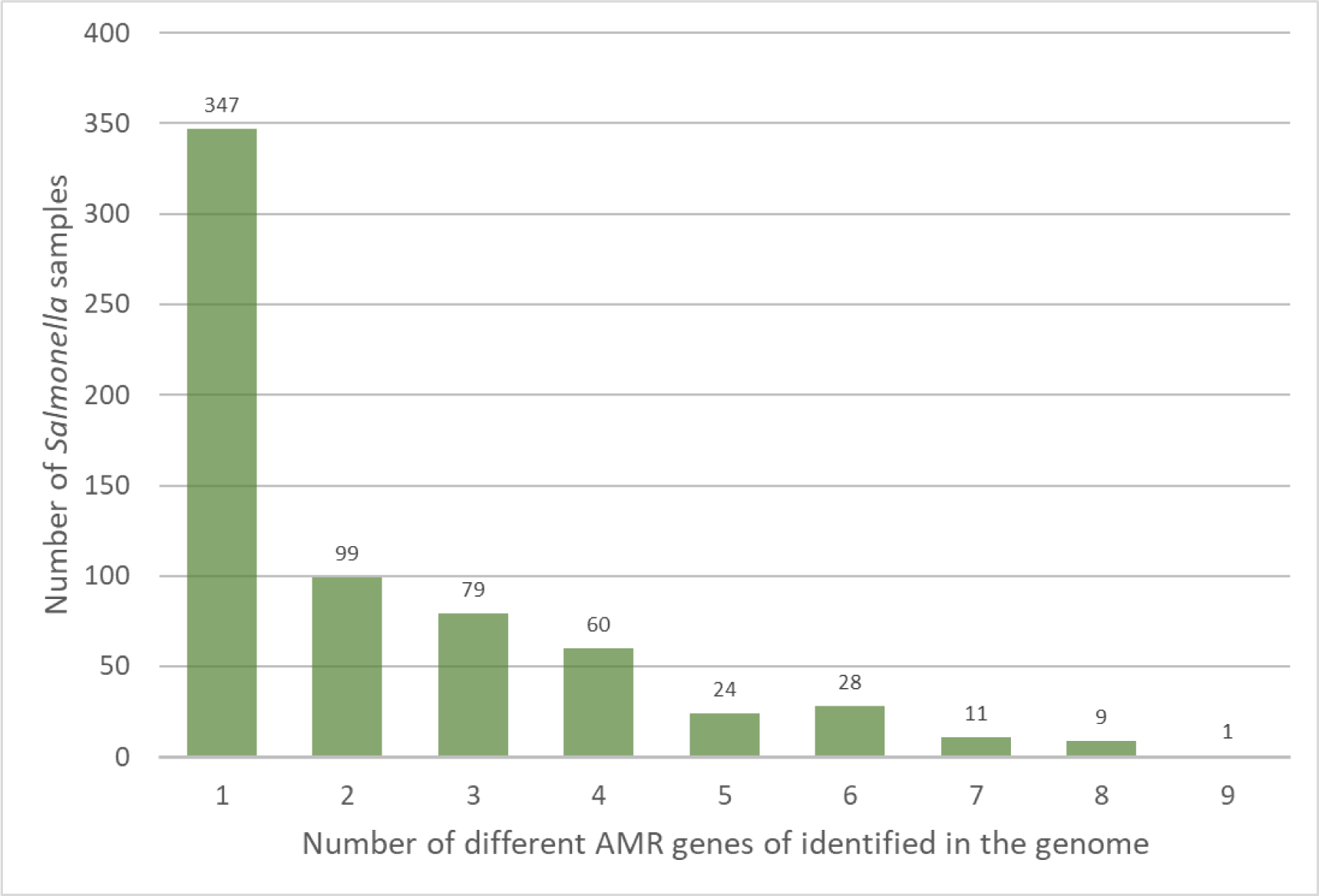
Observed frequency of potential multi-resistance cases found among the *Salmonella* samples sequenced. Number of samples in which from only one to up to 9 different AMR genes have been found.

Another perspective for monitoring antimicrobial resistance is through the surveillance of plasmids harboring these resistance genes. SIEGA flexible tables allow the integration of data to both antimicrobial resistance and plasmid tracking, facilitating a comprehensive analysis. This data can be graphically represented on a phylogenetic tree, which illustrates the STs that have acquired resistance-bearing plasmids. Moreover, this representation can highlight whether such acquisitions have occurred within the same time, geographical region, livestock farm, food processing plant, grocery store or healthcare facility, thus providing critical insights into the patterns and pathways of resistance spread. Figure 5 illustrates a case is the plasmid NZ_AJ437107, which harbors a beta-lactam resistance gene. This plasmid has been acquired in both livestock and food samples with the same sequence type, similar date and from the same geographical location, suggesting a potential selection pressure in certain environments favoring the acquisition of beta-lactam resistance genes.

**Figure 5.**
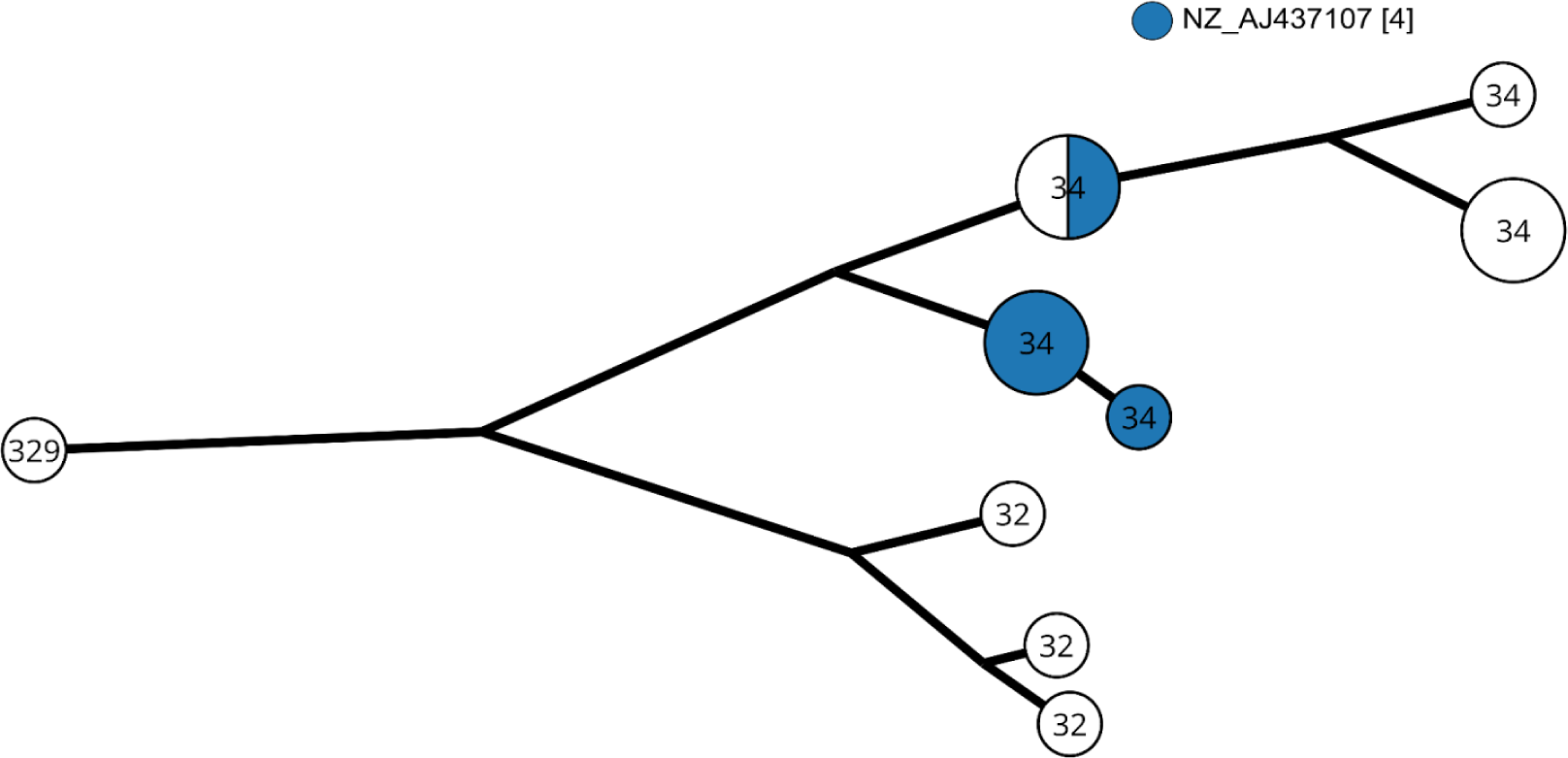
Phylogenetic tree, based on allelic differences (log scale), of Salmonella enterica isolated from the same locality, grouped by allelic profile (the circle size correlates with the number of samples). Blue dots represent strains harboring the plasmid NZ_AJ437107, and the number inside each dot indicates the corresponding MLST for each sample.

### Some successful SIEGA use cases

Despite its incipient use, SIEGA has already proven its usefulness in several cases. Among them it is worth mentioning the investigations carried out in connection with an outbreak of Salmonella Agona, as declared by Norwegian authorities ^36^, entailed, irrespective of field investigations, a comprehensive review of the SIEGA database in pursuit of genomic congruences. This study resulted in the absence of any coincident strains. Another case was the investigation regarding the food alert notification issued under the identifier 2020.5961 within the European Commission Rapid Alert System for Food and Feed (RASFF) ^37^, involving actions that extended beyond on-site measures. These actions included obtaining genomic sequences from food samples supplied by the Finnish food safety authorities. These sequences were then integrated into the SIEGA database. However, no matches were identified both at the time of integration and among subsequent samples added to the system. Further investigations undertaken in relation to the “Joint ECDC-EFSA rapid outbreak assessment” released on July 27, 2023 ^38^, in which Spain was identified as one of the conceivable sources of the suspected transmission vehicle, involved, regardless of field actions, a reassessment of the SIEGA database in search of genomic coincidences, resulting in the non-existence of any matching strain.

These cases clearly illustrate how SIEGA is valuable in ruling out the existence of coincident strains in our database with strains from evaluations of other outbreaks detected at the national or European level, thus facilitating decision-making. On the other hand, alerts are generated that point out possible connections between the stored strains and the ongoing outbreaks, accelerating the possible identification of the source of origin in an outbreak, an aspect that is very difficult to identify with previous methods.

## Discussion

In order to effectively apply the One Health approach to the surveillance and control of antimicrobial resistance, it is crucial to adopt methodologies that ensure consistent results across diverse settings and various sample sources, facilitating comprehensive and reliable data analysis ^39^. Traditionally, surveillance efforts have predominantly centered around specific organisms, prioritizing genotypic and/or phenotypic characteristics that are deemed significant for individual pathogens. Consequently, a multitude of distinct methodologies emerged, giving rise to challenges of consistency, applicability, standardization, and scalability across laboratories and pathogens. Addressing these issues necessitated the implementation of rigorous protocols, regular quality and harmonization controls, as well as the adoption of shared platforms and common controls for result dissemination, thereby mitigating some of the aforementioned concerns ^40^. Nonetheless, the advent of high-throughput sequencing and the accessibility of obtaining comprehensive or nearly comprehensive genome sequences from bacterial isolates within a reasonable timeframe and cost have brought about a profound transformation in the pursuit of surveillance objectives, addressing a significant portion of the previously mentioned challenges. While complete realization remains a work in progress, the trajectory appears unambiguous, with the majority of agencies transitioning toward genome-centric surveillance. This approach presents numerous advantages, notably the capacity to establish connections among surveillance outcomes across diverse levels, locations, and species, which holds particular relevance for the One Health framework ^41,42^.

Currently, the genomic surveillance in Andalusia comprises 6 main pathogens, although new pathogens will be included in brief, like Vibrio or others, pending further decisions. Specifically, *Campylobacter* is one of the major foodborne pathogens of concern in its growing trend of antimicrobial resistance. *C. jejuni* and *E. coli* are the major causative agents, with *C. jejuni* contributing to most of the cases in approximately 90% in the world. Infection is transmitted to humans due to consumption of contaminated food and water ^43^. It is very necessary to establish the chains of transmission between reservoir animals, food and humans. There are few published studies on the genomic characterization of human clinical strains ^44,45^ and their relationship with outbreaks ^46^. It would be very useful to establish sentinel surveillance, such as the one implemented in Ireland recently ^47^. The difficulty in maintaining the viability of *Campylobacter* in environmental samples is a challenge to increase the data that provide comparative information with clinical strains.

Although SIEGA represents a regional solution, Andalusia due to its large, country-like size, can be considered an illustrative case of implementation for a medium-sized region in Europe, serving as an exemplary guide for other authorities wishing to adopt a similar model. Actually, within an international context, SIEGA has yielded results in ruling out the involvement of products originating from Andalusia in outbreaks occurring in other European countries. Moreover, upon reaching a national-level management agreement, it will facilitate the inclusion of non-human origin strains in the European Food Safety Authority (EFSA) sequencing database, as the quality controls and bioinformatics analysis tools described in [reference] have been implemented. This will enable both EFSA and the ECDC at the European level to interact with genomic information originating from Andalusia. It is also important to remark that SIEGA has been designed with the capability to incorporate new domains, such as strains originating from other territories, countries or entities with its multi-level user management system and flexible permissions.

Summarizing, this work presents SIEGA, an advanced integrated One Health system that allows precise surveillance of environmental pathogens, permitting a comprehensive characterization of new isolates with data quality control and data traceability included. SIEGA facilitates automated generation of customized reports that include similarities with other samples, represented as a dendrogram, potential AMR or virulence genes detected, among other details. The alerting functionality of SIEGA, that alerts as immediately as one sample is introduced that meets a predefined similarity conditions, is a revolutionary tool for detecting transmission chains at an early stage. In addition, the possibility of crossing metadata and representing them over sample dendrograms allows performing detailed retrospective studies of different sample features (e.g., emergence or transmission of AMR, etc.) Moreover, since SIEGA connects environmental with clinical samples it allows tracing clinical occurrences back to their environmental origins, permitting rapid interventions targeted to the source of the outbreak. In addition, the success in discarding relationships in the use cases underscore the intricate challenges in elucidating the etiology and dynamics of microorganism-related incidents. The pursuit of genomic concordance within the SIEGA database, although yielding no direct matches, serves to illuminate the genetic diversification that these pathogens can exhibit. These episodes emphasize the need for continued vigilance, inter-agency cooperation, and cutting-edge molecular methodologies to fortify our comprehension and management of such microbial phenomena.

Thus, by leveraging high-throughput sequencing technologies, advanced bioinformatics tools, and robust data sharing platform, SIEGA offers a comprehensive view of the genetic landscape of pathogens across different geographical regions and host populations.

## Conclusions

Centralized circuits of genomic surveillance based on whole genome sequencing (WGS) as SIEGA provide a convenient and efficient way to monitor infectious diseases and detect outbreaks in real-time ^48^. Such facilities enable the characterization and relatedness determination of bacterial isolates, aiding in tracking transmission patterns and implementing effective infection control measures ^4^. Moreover, WGS has been incorporated into public health surveillance systems, offering significant contributions to outbreak investigations, infection prevention, and control ^48^. The potential integration of WGS into epidemiological investigations has been highlighted, emphasizing the need to establish optimal models for data integration and evaluate public health impacts resulting from genomic surveillance ^49^. By harnessing the power of whole-genome sequencing, centralized circuits of genomic surveillance offer immense potential for improving disease surveillance and response, ultimately contributing to the overarching goals of the One Health framework ^1,4,39^.

## Methods

### DNA extraction protocol

For the majority of the samples, DNA extraction was conducted using the PureLink Genomic DNA Mini Kit (Invitrogen). Subsequently, quantification of the extracted DNA was performed using the QUBIT FLEX fluorometer, and the quality of the eluate was assessed through electrophoresis using the Egel Power Snap Electrophoresis Device.

### Sequencing

The majority of the samples have been sequenced at the CABIMER Genomic Unit, with a significant contribution from Listeria sequences obtained from the National Center of Microbiology (469 samples). These sequences were incorporated under a mutual agreement for sequence exchange established between the Andalusian Public Foundation Progress and Health-FPS and the National Center of Microbiology in 2020.

Whole genome sequencing of the isolates has been carried out at the CABIMER genomic facility. Library construction and sequencing was performed at Genomics Core Facility of CABIMER. DNA libraries were prepared using Nextera DNA Flex Library prep kit (Illumina) following the manufacturer’s instructions. High-throughput sequencing was performed on the NextSeq 500 Sequencing System (Illumina).

### Sequencing data processing

The raw reads are filtered using the fastp ^50^ application, followed by a search for potential contamination from other organisms using the kraken2 ^51^ and bracken ^52^ applications. Coverage quality control is carried out using qualimap2 ^53^. Subsequently, with quality-controlled reads, a de novo assembly is performed using SPAdes ^54^ and the quality of the assembly is assessed using QUAST^55^.

### Sample typing

MLST (Multi-Locus Sequence Typing) is acquired using the MentaLiST ^56^ application for those organisms whose databases are updated within this tool. CgMLST (core genome Multi-Locus Sequence Typing) profiles are generated using Chewbbaca ^57^ with the schema available in the Chewie Nomenclature Server (chewie-NS) ^58^ for *Listeria monocytogenes. Salmonella enterica* and *Escherichia coli* schemas are filtered based on the EFSA gene list ^59^ CgMLST profiles. Moreover cgMLST profiles based in other databases are obtained using a custom script that utilizes the BLAST+ ^60^ tool and allelic profiles from the following public databases as references: Pasteur Institute ^61^ for *Listeria monocytogenes*, EnteroBase ^33^ for *Yersinia enterocolitica, Salmonella enterica* and *Escherichia coli* and PubMLST ^62^ for *Campylobacter jejuni/coli.* Additionally, the applications LisSero ^63,64^, SeqSero2 ^65^, and SerotypeFinder ^66^ are employed to determine the serotype of *Listeria monocytogenes*, *Salmonella enterica*, and *Escherichia coli*, respectively.

### AMR genes

To identify antimicrobial resistances, two approaches were employed. Firstly, the ResFinder ^67^ application was utilized, along with its dedicated database. Secondly, the ABRicate ^30^ tool was employed in conjunction with various databases, namely CARD ^32^, MegaRES ^68^, and ARG-ANNOT ^69^.

### Virulence genes

To identify virulence genes, two applications were employed: VirulenceFinder ^70^, with its dedicated database and ABRicate using the Virulence Factor Database (VFDB) ^31^ as a reference.

### Plasmids

To identify plasmids present in the samples two approaches are employed: the PlasmidFinder ^71^ application and the Mash Screen ^72^ application combined with the PlasDB database ^73^.

### Core genome determination and phylogenetic analysis

Using the parSNP ^74^ application, the core genome is analyzed, SNP selection is performed, and subsequent phylogenetic trees are constructed based on SNP differences for each of the organisms present in SIEGA. On the other hand, allelic differences are obtained using a custom script that compares the cgMLST of each sample for each organism, generating a matrix for each organism. This matrix is then utilized by GrapeTree ^27^ to construct the phylogenetic tree based on allelic distances, which could be visualized GrapeTree or Taxonium ^28^. The ETE3 ^75^ application is used for the selection and generation of sub-trees included in the reports. The reference sequences (NCBI database identifiers) used for the different species were for *Listeria monocytogenes* NC_003210.1, *Salmonella enterica* NZ_SRHS01000001.1, *Escherichia coli* NC_000913.3, *Campylobacter jejuni / coli* NZ_CYQA01000001.1, *Yersinia enterocolitica* NZ_CQAE01000001.1 and *Legionella pneumophila* NZ_CP015941.1.

## List of abbreviations

AMR: Anti-Microbial Resistance
CC: clonal complexes
cgMLST: core genome Multi-Locus Sequence Typing
CSV: Comma-separated values
ECDC: European Centre for Disease Prevention and Control
EFSA: European Food Safety Authority
LIMS: Laboratory Information Management System
MLST: multi-locus sequence-based typing
RASFF: Rapid Alert System for Food and Feed
SIEGA: Sistema Integrado de Epidemiologia Genomica de Andalucia
SNP: Single Nucleotide Polymorphism
ST: Sequence Types
SVEA: Sistema de Vigilancia Epidemiológica de Andalucía
WGS: Whole genome sequencing

## Acknowledgements

This work was supported by the Instituto de Salud Carlos III (ISCIII), co-funded with European Regional Development Funds (ERDF) (grant IMP/00019), it has also been funded by Consejería de Salud y Consumo, Junta de Andalucía (grants COVID-0012-2020, PIP-0087-2021), and by grant ELIXIR-CONVERGE - Connect and align ELIXIR Nodes to deliver sustainable FAIR lifescience data management services (AMD-871075-16), funded by EU – H2020. CSCS was funded by a Juan de la Cierva grant (FJC2021-046546-I) from Ministerio de Ciencia e Innovación.

